# Causal analyses, statistical efficiency and phenotypic precision through Recall-by-Genotype study design

**DOI:** 10.1101/124586

**Authors:** Laura J. Corbin, Vanessa Y. Tan, David A. Hughes, Kaitlin H. Wade, Dirk S. Paul, Katherine E. Tansey, Frances Butcher, Frank Dudbridge, Joanna M. Howson, Momodou W. Jallow, Catherine John, Nathalie Kingston, Cecilia M. Lindgren, Michael O’Donavan, Steve O’Rahilly, Michael J. Owen, Colin N.A. Palmer, Ewan R. Pearson, Robert A. Scott, David A. van Heel, John Whittaker, Tim Frayling, Martin D. Tobin, Louise V. Wain, David M. Evans, Fredrik Karpe, Mark I. McCarthy, John Danesh, Paul W. Franks, Nicholas J. Timpson

## Abstract

Genome-wide association studies have been useful in identifying common genetic variants related to a variety of complex traits and diseases; however, they are often limited in their ability to inform about underlying biology. Whilst bioinformatics analyses, studies of cells, animal models and applied genetic epidemiology have provided some understanding of genetic associations or causal pathways, there is a need for new genetic studies that elucidate causal relationships and mechanisms in a cost-effective, precise and statistically efficient fashion. We discuss the motivation for and the characteristics of the Recall-by-Genotype (RbG) study design, an approach that enables genotype-directed deep-phenotyping and improvement in drawing causal inferences. Specifically, we present RbG designs using single and multiple variants and discuss the inferential properties, analytical approaches and applications of both. We consider the efficiency of the RbG approach, the likely value of RbG studies for the causal investigation of disease aetiology and the practicalities of incorporating genotypic data into population studies in the context of the RbG study design. Finally, we provide a catalogue of the UK-based resources for such studies, an online tool to aid the design of new RbG studies and discuss future developments of this approach.

## Introduction

### Bottom-up genetics and its application to epidemiology and mechanistic understanding

Genome-wide association studies (GWAS) have identified thousands of common genetic variants related to complex traits and diseases^1^. More recently, studies with sequencing data have extended these discoveries to less common genetic variation^2-7^. However, while these studies can detect new associations of genetic variation with a variety of complex traits and open new avenues for exploring underlying biology, they typically provide limited information about function or mechanism. Bioinformatic analyses, studies using appropriate cell lines or animal models and applied genetic epidemiology have all been used to meet the problem of understanding such genetic associations and dissecting causal pathways; however, what is less often seen is the use of genetic data in the design of new studies aimed at elucidating associations and causal relationships. We describe the motivation for and characteristics of genotype-directed deep phenotyping studies -recall-by-genotype (RbG) - and why they can be useful in this context. We consider the efficiency of this approach and the likely value of RbG studies for the investigation of disease aetiology and pathophysiology. We discuss the practicalities of incorporating genotypic data into population-based study designs and provide a catalogue of UK-based study resources and an online tool to aid the design of new RbG experiments.

### Rationale for genotype-based sampling strategies

The collection of informative and detailed phenotypic measurements is needed to help elucidate pathophysiology and disease aetiology, but obtaining data needed in the populations required can be difficult and expensive^8^. Often, the complications of costs and availability render ideal phenotype assessment impractical and lead to situations where measurement precision/quality or proximity to underlying biology is compromised by the use of cheaper pragmatic approaches. These can be entirely suited to reveal crude associations, but often require further work in order to allow a deeper understanding of the original signal or use of it in an epidemiological context. A good example is that of genetic contributions to the common paediatric disorder, asthma. Whilst the known heterogeneity of this important health outcome presents an opportunity to use genetics and detailed phenotyping as a tool for dissection, current studies have employed broadly defined phenotypes (i.e., doctor diagnosed asthma, persistent or troublesome wheeze) in an effort to obtain sufficient sample sizes to investigate this complex trait^9^; ^10^. This approach has allowed for the discovery of reliable associations and flagged important biological pathways involved in disease, but has prompted the study of precise intermediate phenotypes to inform mechanism or gain resolution^11^; ^12^. Indeed, the desire for measurement of informative “endophenotypes” has been called for elsewhere in an effort to try and discover the otherwise missing intermediate steps between genotype and complex health outcome^13^.

To address this, targeted studies can be undertaken that allow the examination of dense phenotypic information in sample sizes that are both financially and practically feasible and have the potential to optimise analytical power by sampling in an informed manner. Studies that recruit subgroups of participants from the extremes of phenotype distributions (such as lean and obese individuals) have been used in epidemiological investigations for many years; however, these studies suffer well known limitations of observational epidemiology^14^. In contrast to these, RbG studies use naturally occurring genetic variants robustly associated with specific traits and diseases to stratify individuals into groups for comparison and are novel and beneficial for two reasons. Firstly, by exploiting the key properties of genetic variants that arise from the random allocation of alleles at conception (Mendelian randomization (MR))^14-16^, RbG studies enhance the ability to draw causal inferences in population-based studies and minimise problems faced by observational studies (**Figure 1**)^14^ ^17^. Secondly, focusing phenotypic assessments on carefully selected population subgroups can improve insight into mechanism and the aetiology of health outcomes in a cost-efficient manner through targeted deployment of more precise and informative phenotyping across already known biological gradients. Together, these features can help dissect existing genetic associations and make efficient use of the genetic prediction of risk factor exposure through the execution of novel and genotype-informed studies.

**Figure 1.**
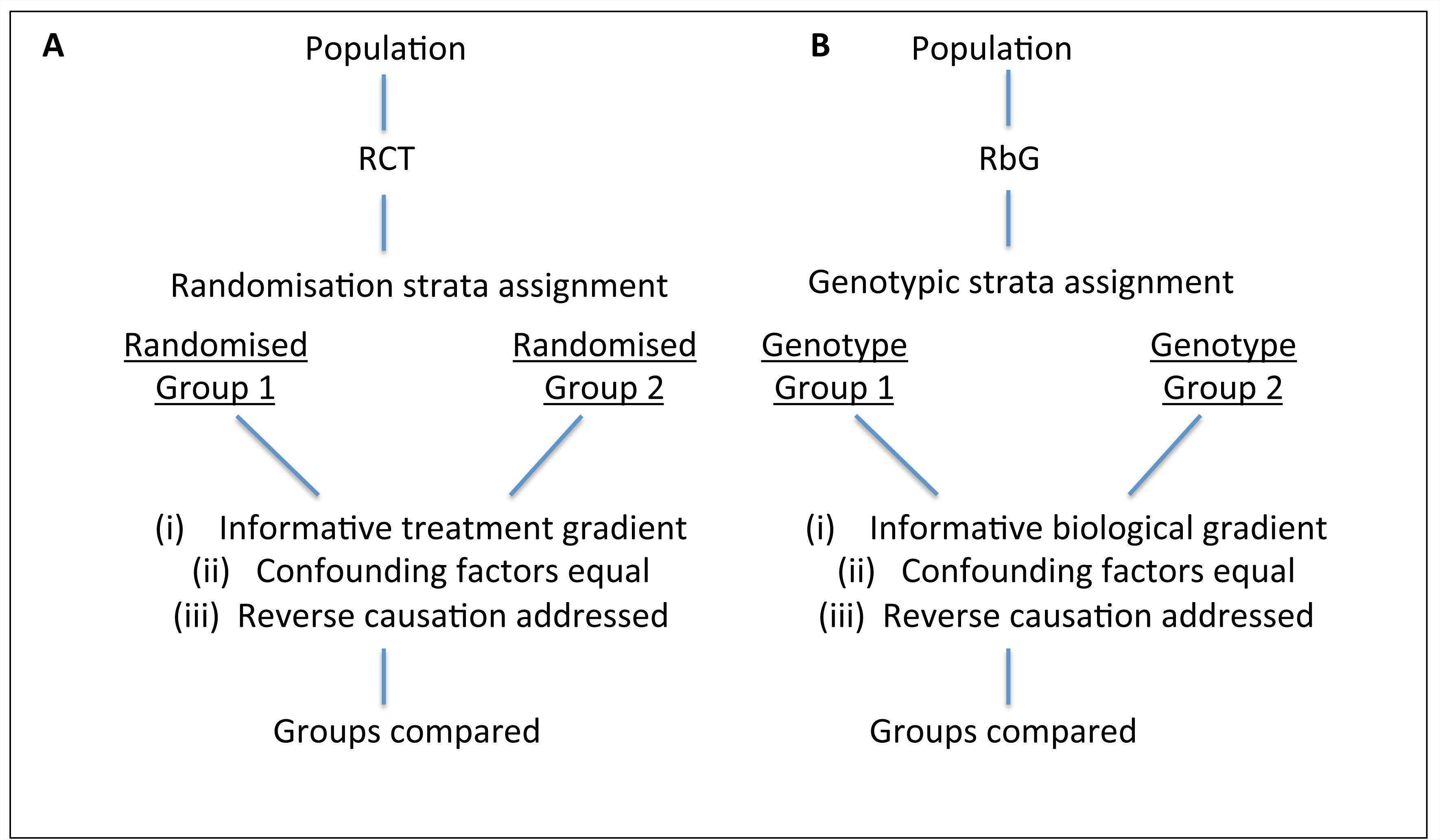
Properties of RbG strata compared to randomised control trials. (A) For randomised controlled trials (RCTs), participants are randomly allocated to intervention or control groups. Randomisation should equally distribute any confounding variables between the two groups. For Recall-by-Genotype (RbG) studies, strata used to define groups are “randomised” with respect to genotypes and, analogous to RCTs, potential confounding factors are equally distributed between groups. Hence, RbG studies are not subject to reverse causality or confounding factors with respect to the phenotype under study by the stratifying genotype.

### Study design, applications and considerations

Forms of RbG have appeared in designs looking to optimise RCT and investigate pharmacogenetic relationships^18-22^, but have not been fully described for population-based resources. Indeed, RbG study designs are likely to develop further and below we present design considerations for RbG in simple form, splitting the approach into two categories for the purpose of description; RbG using a single variant (RbG^sv^) and RbG using multiple variants (RbG^mv^). These have the same inferential properties based on the properties of genetic data (above); however, they describe differing analytical scenarios and point to the potential variety in this application of human genetics.

**RbG^sv^** studies can be considered the most intuitive type of RbG, whereby strata defined by a single genetic variant are used as a basis for the recall of samples or participants for further phenotypic examination. This type of RbG study may focus on functional variants known to induce a direct biological change; however, genetic variants may also be chosen if they have uncharacterised or predicted effects (i.e., loss-of-function variants, cis-regulatory variants or intronic variants that alter DNA-protein binding at potential drug targets)^23^. These variants provide natural experiments able to yield information about the specific role of biological pathways as well as gradients within them and potentially inform on both the safety and efficacy of medicines. For RbG^sv^ studies, participants or patients or their samples are recruited and phenotypes measured based on genotypic groups in a manner not dissimilar to the arms of a clinical trial. Recall in this way yields groups in which detailed phenotyping can be undertaken in order to assess the specific impact of a genetic change or the aetiology of an outcome. An early example of this approach was an investigation of the effects of the peroxisome proliferator-activated receptor-γ (PPARγ) Pro12Ala polymorphism on adipose tissue non-essential fatty acid metabolism^24^. Key examples of RbG^sv^ are included in **BOX 1** and there are studies currently underway that have had protocols reported in advance of their completion^25^; ^26^.

The precise sampling strategy for RbG^sv^ will depend on properties of the target variant and predictions about its mode of inheritance. Here we consider the implications of recruiting an equal number of major and minor homozygotes (or carriers of the minor allele (heterozygotes) if frequency is very low) in an effort to maximise available biological contrast. However, if it is known, consideration of the appropriate genetic model can aid design (particularly where effects are dominant) and an alternative strategy is to recruit equal numbers of all three genotype groups^27^. For homozygote sampling, there exists an optimal range of minor allele frequency (MAF) at which RbG^sv^ studies outperform random recall designs and the power of RbG studies increases beyond random recruitment particularly at low (MAF< 10%) to very low (MAF< 1%) allele frequencies (**Figure 2A**). This efficiency, however, comes at some cost as recruiting sufficient participants or samples with low or very low frequency genotypes requires much larger bioresources (with genetic information) from which to recruit than for high frequency genotypes. For instance, consider a study recruiting individuals based on a genetic variant with a MAF of 5% and requiring a total sample size of 50. The genotyped bioresource sample would need to contain at least 10,000 individuals in order to identify 25 minor allele homozygotes (assuming Hardy-Weinberg equilibrium) (**Figure 2B**). Given that not all of these participants will be eligible or willing to participate in the RbG study (or where alternative group size ratios are required), the required bioresource sample size is likely to be even larger.

**Figure 2.**
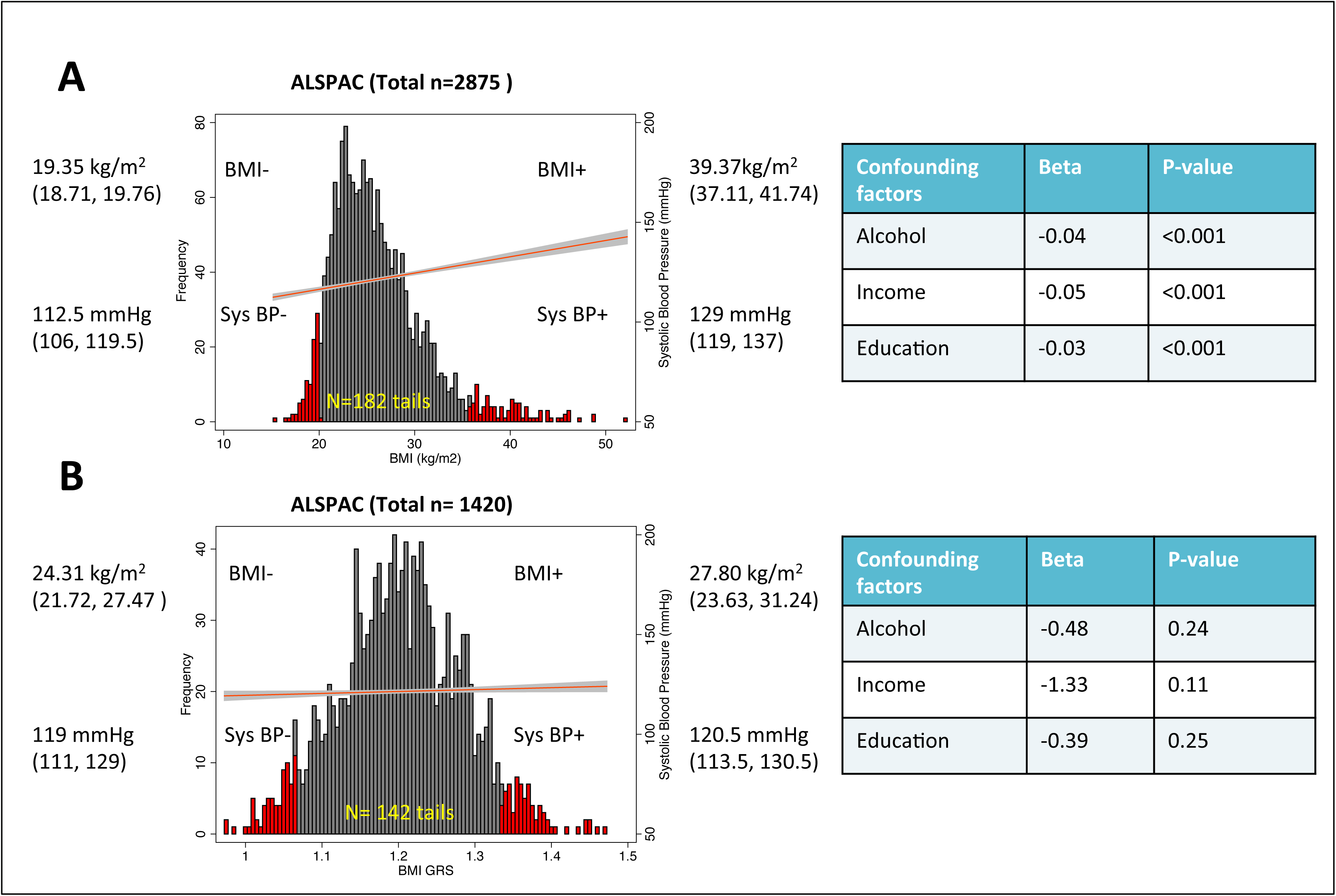
Comparative power: RbG^sv^ versus random recall study designs. For the purpose of the power calculations presented, it is assumed that the RbG^sv^ study will recruit an equal number of major and minor homozygotes, and that the Type I error rate (alpha) is 0.05. (A) Top panel: A direct comparison of power (y-axis) achieved by an RbG^sv^ study design (calculated using a basic two-tailed t-test) versus a random sample selection (calculated using a X test assuming an additive genetic model) design for a given minor allele frequency (MAF) and standardised effect size. The x-axis is the total sample size of the recall experiment. Lower panel: A representation of the difference (y-axis) between the power within an RbG^sv^ study design and that from the equivalent random recall experiment. (B) A representation of the trade-off when undertaking an RbG^sv^ experiment in the expected number of participants with genotypic data (y-axis) needed in order to recruit sufficient minor homozygotes for a given RbG^sv^ study sample size (x-axis) and MAF (assuming a 100% participation rate).

**RbG^mv^** designs can be appropriate when interest is focused on the impact of an exposure and whilst collections of genetic variants can be used to predict biological mechanisms or pathway activity^28^, risk factor exposure will be the focus of these study designs. Rather than employing a single variant, a genetic proxy for exposure is generated by combining genotypes that explain some proportion of the variance in the exposure of interest. The choice of genetic variants for RbG^mv^ studies can be complex and relies on the ability of genotypic variation to act as a reliable proxy measure for the exposure of interest. Genetic variants associated with complex traits often explain only a small proportion of variance in that trait and a strategy employed to try and recover some of the consequent lack of power of single variant analyses is to generate genetic risk scores (GRSs), or aggregate scores of genetic difference in traits of interest^29-31^. The use of multiple genetic variants in this way can increase the precision of the causal estimate compared with those derived using separate genetic variants^32^. Once a GRS is constructed within the study sample set targeted for RbG (usually as the sum of allele dosages at risk variants weighted by their beta coefficients obtained from an independent GWAS for the exposure of interest), individuals can be ranked on the basis of this score. This summary of phenotypic difference can then be used to stratify participants for recall (**Figure 3**). Actual selection of individuals from extremes of the GRS will be dependent on the number and frequency of the variants forming the score, their effect and the number of participants (or samples) available. In addition, the actual score carried by specific individuals recruited into the study will differ in terms of the contributing genetic variants and causal mechanisms by which these variants impact the trait of interest; however, differences across the genetic stratum will carry the same inferential properties as RbG^sv^. An example of an RbG^mv^ study is included in **BOX 1.**

**Figure 3.**
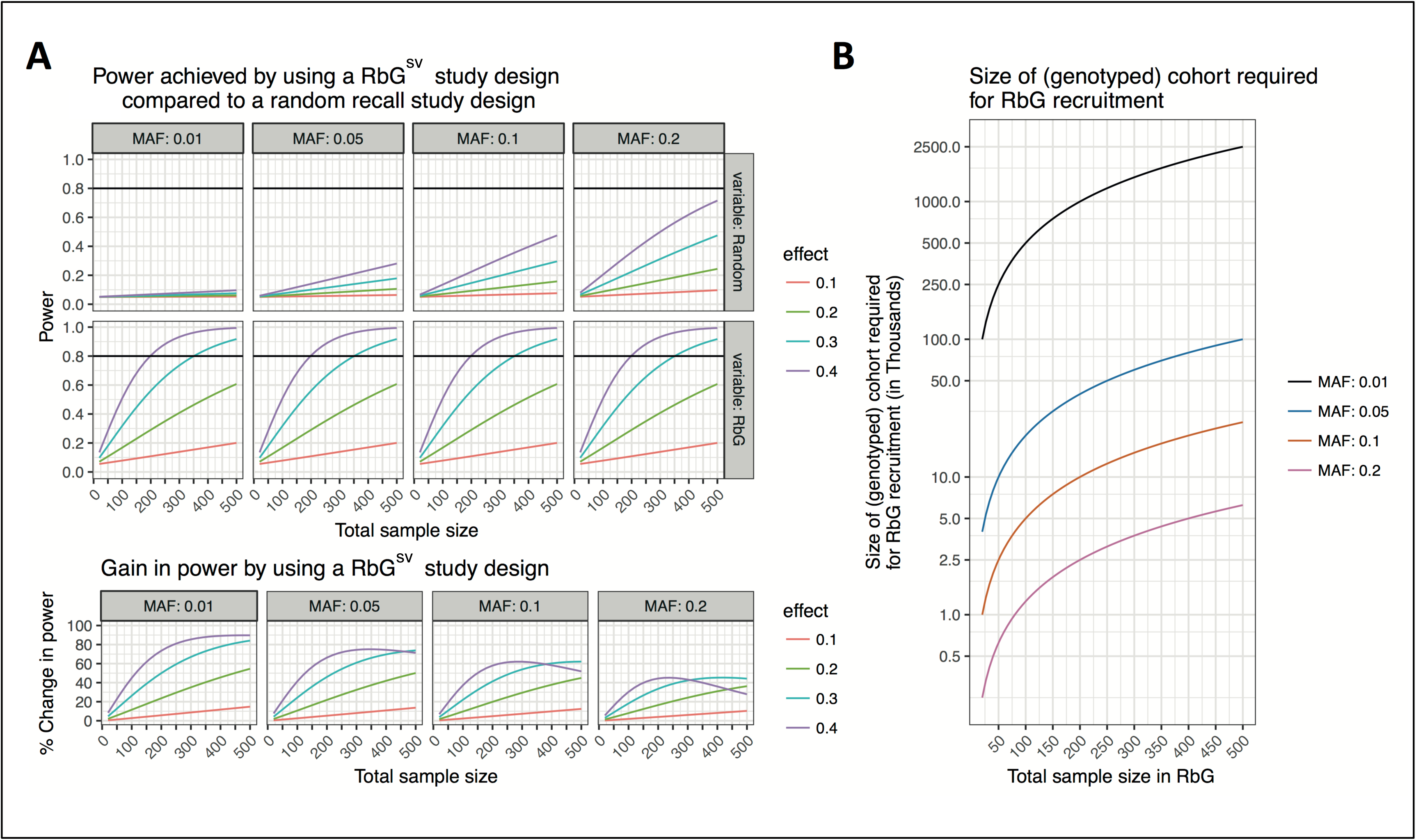
Contrast between phenotype and genotype-based sampling strategies. The dark grey histograms show the distributions of (A) body mass index (BMI) using an extreme-phenotype recall study design and (B) the BMI genetic risk score (GRS) using a Recall-by-Genotype (RbG) study design in the Avon Longitudinal Study of Parents and Children (ALSPAC). Red bars represent the top and bottom 30% of these distributions. Mean differences in BMI, SBP and confounding factors (alcohol, income and education) were compared between the top and bottom 30% of the (A) BMI and (B) BMI GRS distribution. (A) For extreme-phenotype recall studies, participants at the extreme ends of the phenotypic distribution are invited to participate in the study. As an exemplar of this, phenotype data from 2,875 individuals in ALSPAC was used (for full details see supplementary material). Whilst differences in BMI and SBP are observed between the top and bottom 30% of the BMI distribution, extreme-phenotype sampling strategies are often prone to confounding and reverse causality (as shown by the association of the recalled strata with confounding factors). (B) In contrast, RbG studies have the ability to generate reliable gradients of biological difference in combination with essentially randomised groups. As an exemplar of this, genetic data from 1,420 individuals in ALSPAC was used to generate a BMI genetic risk score (GRS) (for full details see supplementary material). Differences in BMI and SBP are observed between the top and bottom 30% of the BMI GRS distribution that are not prone to confounding factors and reverse causality (as shown by the lack of association of the recalled strata with confounding factors).

For RbG^mv^ studies, the greatest gain in power occurs when the sample groups are recruited from the most extreme part of a GRS distribution (to yield an exposure contrast of interest) but, where sufficient numbers of participants (or samples) are available given the new phenotype to be measured. This type of sampling is a function of the genetic architecture of a given exposure (the nature of genetic contributions to a trait) and not just the variance explained. In order to design this type of GRS-driven RbG experiment, firstly the GRS needs to be modelled and the number of individuals eligible for recruitment in any given sample estimated given the properties of the GRS used. Unlike RbG^sv^ designs, where the frequency of one variant determines the sampling frame, the frequency and effects of a collection of implicated loci determine the number of participants available.

### Statistical power

Key to the design toolbox for RbG studies is an appropriate framework for the assessment of statistical power. Undertaken correctly, power calculations illustrate the optimum conditions in which one would consider using an RbG experiment as an approach as opposed to more conventional sampling methods. As described above, power for RbG^sv^ studies can be calculated based on the proposed sample size and the balance of major homozygotes to minor homozygotes/heterozygotes therein (this can be adjusted to optimise power as in a conventional case-control design), the phenotypic properties of the outcome of interest (specifically, the standard deviation) and the anticipated difference in outcome by recall group. In this simple form, RbG^sv^ studies recruit individuals (or samples) into two groups that are independent of allele frequency and thus power can be estimated assuming a standard two-sample two-tailed t-test for the difference in means.

Power for RbG^mv^ studies can be considered as a two-part process reflecting not only the properties of the outcome measure, but of the exposure gradient being measured in proxy by the genotype or GRS in question. This can be modelled using properties of the genetic variants and their aggregate effects to predict (i) the distribution of the GRS, (ii) the number of participants in the tails of the GRS for any given sample size and (iii) the magnitude of the association between set thresholds of the GRS and the exposure of interest. Given a satisfactory exposure gradient has been confirmed for the GRS in question, the second part of the process follows that of RbG^sv^ studies (i.e., the basic consideration of a recall sample to detect biologically informative differences in the outcome phenotype between GRS predicted exposure groups). At this point, the anticipated relationship between the exposure phenotype and the proposed outcome measure (although not likely to be known precisely) will be important in determining the relative gain of the study over a random sampling design. Consideration for both the practicalities of sampling at the tails of a GRS (i.e., that one cannot know how many participants who are invited to participate will eventually complete the study) and the nature of the GRS distribution (especially if composed of very few genetic variants) should be made.

For both RbG^sv^ and RbG^mv^ approaches, there may be situations where power can further be enhanced (and biological effect clarified) when comparing genotype-driven recall groups also group- or pair-matched for characteristics such as age, sex and body mass index (BMI). Whilst access to larger sample sizes may reduce the need for matching to preserve power, matching may also be advantageous when there are genotype-driven differences in the potential for ascertainment (e.g., for an early-onset fatal disease, or in selecting non-diabetic individuals for a study of a diabetes risk variant) and this approach has been exercised in existing studies^23^; ^33^. There is, however, a danger that such manoeuvres can exacerbate the potential for particular types of study bias^34^ and the pros and cons of these decisions need to be weighed carefully in study design.

To facilitate the design of RbG experiments based on the scenarios outlined above as “RbG^sv^” and “RbG^mv^”, we have prepared an online tool for guiding researchers through these steps that is available at the Recall-by-Genotype Study Planner. The methods used to calculate power for RbG studies are described in more detail in the supplementary material. Current methods do not incorporate matching of RbG groups and consequently are likely to be conservative.

### Ethical considerations when utilizing genetic markers in recall studies

RbG is a potentially powerful research design, but it creates ethical challenges. The RbG approach is inextricably linked to the issue of disclosing potentially sensitive individual results^35^; ^36^ and places an emphasis on transparency and communication with participants. Suggestions for general approaches for making decisions about what (if any) information to disclose to study participants have been framed around four fundamental ethical principles of research: beneficence, respect, reciprocity and justice^37^. Despite this, there is little published academic work regarding the specific ethical issues in RbG studies. A small body of literature suggests a need for “bottom-up research” to be monitored by an independent governance body^38^ and that the issues presented to RbG studies are not new but common to those faced by other approaches, such as the use of medical records^39^. Qualitative data that does exist around this topic compared the experiences of patients (those with the disease of interest) to those of “healthy volunteers” (recalled from a biobank) following their recruitment on the basis of genotype^40^. This research has found that whilst patients expressed “no concerns” about the eligibility criteria, “healthy volunteers” did not always comprehend the study design or why they had been chosen. This led in some cases to participants assuming a degree of meaningfulness to the genetic data that was unwarranted but nevertheless caused them to feel anxious. Indeed, work explicitly interviewing participants about their opinions and response to RbG methods has explicitly recognized the feelings of trust present in groups of engaged individuals and thus the responsibility placed onto researchers for the handling of potentially sensitive and disclosive studies^41^.

The very nature of RbG designs highlight a central tension between avoiding the possibility of participant harm through revealing unwanted or misunderstood information and being open and clear when explaining how and why participants are being recruited into studies^35^;36. In healthy volunteers, it is unlikely that the genetic information used for recruitment to most RbG studies will be either immediately clinically valid or useful, as the precise function of the genetic characteristics will presumably be unknown. However, this does not diminish the need to clearly communicate the study protocol to participants and why they, specifically, have been recruited. It is of course possible to envisage a situation whereby the threshold of clinical utility obtained through an RbG study is not reached but the genetic information could still be of interest to the participant. This is most likely to occur when the focus of the RbG study is a rare variant with a relatively large effect on a common trait or disorder. The employment of sensible mechanisms for assessment of data quality and routes for appropriate feedback (as considered in detail for sequencing studies elsewhere)^42^ will clearly be the accepted mode for RbG studies with large effects. However, the issue of addressing a specific genotype-driven effect does serve to illustrate a key advantage of RbG studies over less hypothesis-driven genomic research. In this case, it is potentially easier to anticipate the nature of findings for a given recall stratum and therefore the potential relevance of those findings to participants^35^; ^36^.

Finally, it is important to note that as RbG experiments and study samples in their own right become more common, specific permission and agreement to allow appropriate feedback at the time of recall is being incorporated into initial resource design. Issues of selection, disclosure and the dissemination of results will likely continue to raise challenges, but the nature of participant response to genetic data use in this way seems likely to follow that when faced with more conventional study designs.

### Resources

Despite potential advantages of genotype-based sampling strategies, they have so far been underutilised, partly because limited infrastructure has existed to support them. However, at a time where the potential value of population-based human genetics is being realised in a clinical context^43^, recent developments have changed the scientific landscape. A growing number of bioresources have been established or re-purposed to enable RbG studies (patient and population-based studies table, **Table 1**) and are ready for coordinated deployment to maximise RbG designs. A second factor has been the continued fall in genotyping and sequencing costs, which has accelerated discovery and enabled genetic characterisation of large cohorts consented for RbG studies. Finally, in recent years a number of RbG studies with important findings have been reported that highlight the value of the approach and illustrate key variations on it.

**Table 1.**
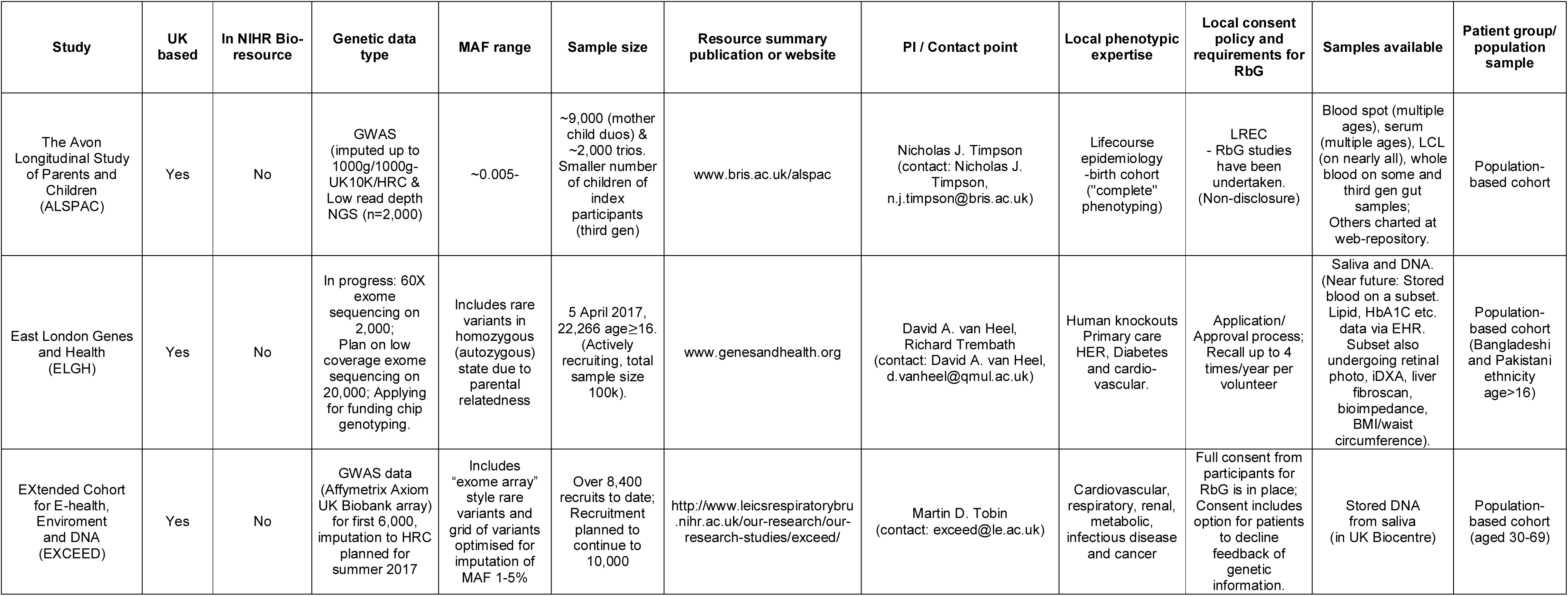
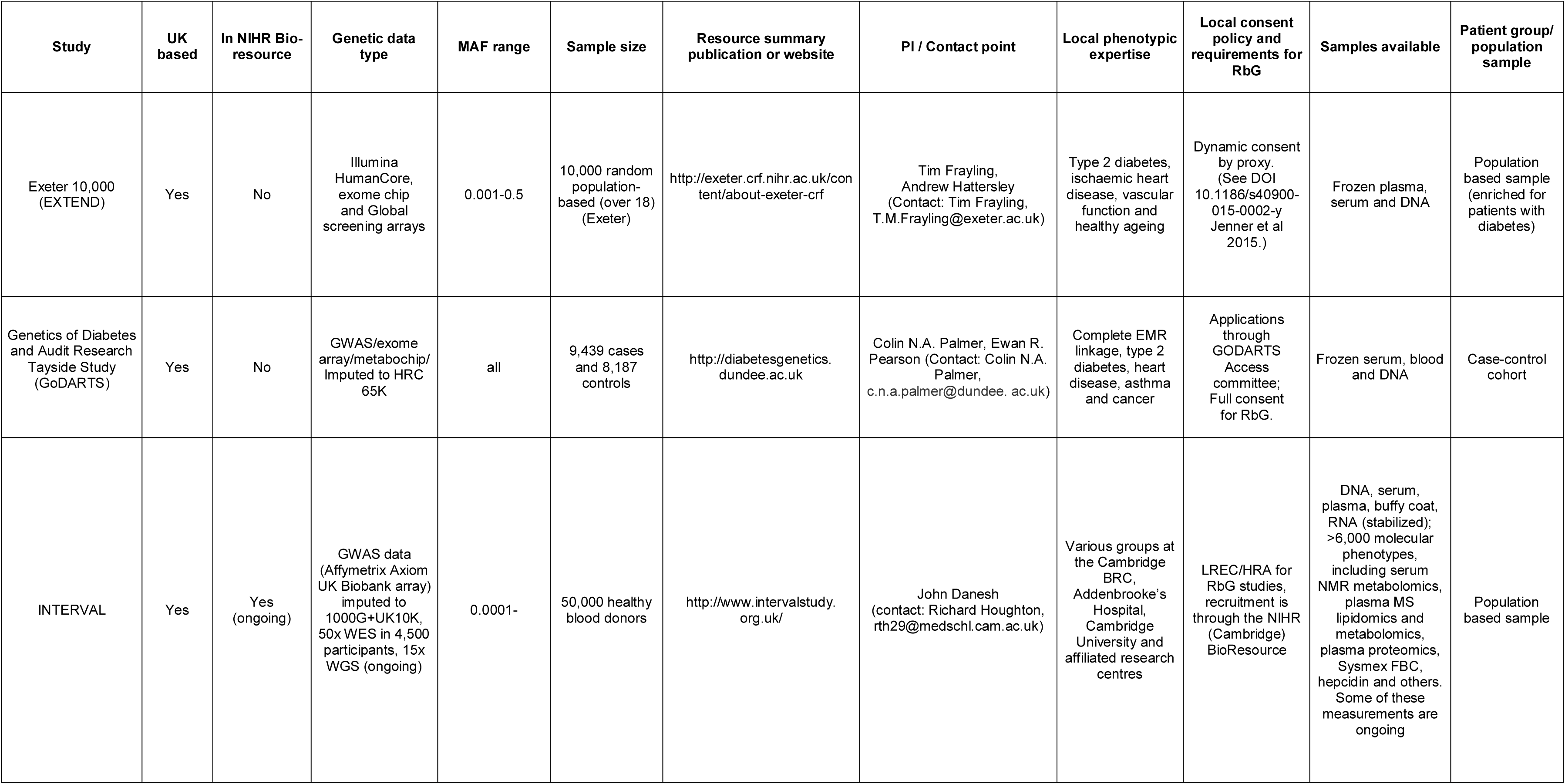
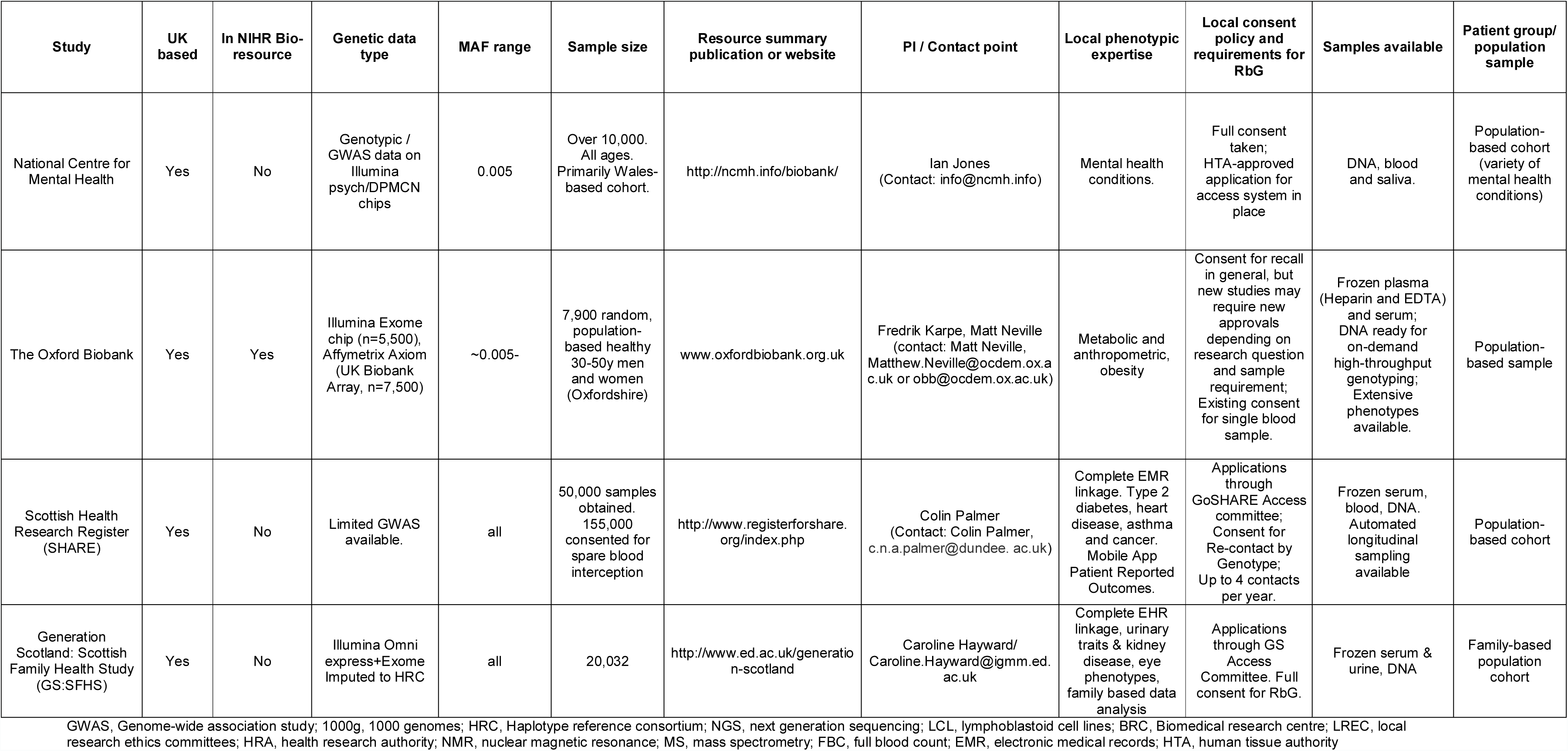
UK Patient and population-based studies available for RbG studies.

### Future directions and developments

RbG studies take an efficient approach to engineering enhanced representation of informative biological gradients found within very large population/participant-based collections. This is not a novel approach to epidemiological studies, where phenotype-driven selection has been a mainstay for the purposes of maximizing analytical power. The novelty here comes from the selection process being based on genetic strata, which not only have the ability to recapitulate the biological pathway changes or exposure differences desired but to do so using reliably measured, reproducible and randomly allocated markers. In the correct conditions, this approach has the potential to be both cost-effective and biologically informative. This positions RbG as potentially a key contributor to the dissection and translation of genetic associations that are being delivered from genome-wide approaches.

In order for the RbG approach to prove of greatest benefit, large-scale population and patient-based records of genotypic variation data with appropriate consent are needed. The ability to measure genetic variation reliably (including those with low MAFs) is an important asset to this approach and has been facilitated by the swathe of both GWAS analyses and imputation development that has occurred over the last 5-10 years. To take this further, the presence and development of effective networks of RbG-ready collections will undoubtedly be required. Not only will these networks allow for the look-up and access of rare variant carriers in reasonable numbers, but the nodal bases of phenotypic expertise will help to develop and exercise the real value of RbG studies in deep phenotyping and enhanced statistical power.

This paper has attempted to advance RbG as an innovative and potentially valuable study design in its simplest form within population-based studies. There are, however, specific adaptations and potential limitations that are relevant to this approach. On the assessment of power, the development of current approaches to simplified RbG conditions provide conservative estimates of the performance of RbG studies, but should be developed to formally incorporate the application of group and pair-based matching. As undertaken in RCTs, this approach has the potential to increase statistical efficiency further and to go beyond the randomisation attributed solely to genotypic group allocation, especially in small sample sizes and where biases may be present. In addition to this, it is equally important to consider the potential of employing variants of specific functional effect or a set of genetic variants^44^ that act together, interact or are responsible for specific pathway effects. With increased information about the gravity of specific and functional genetic changes and a growing collection of whole genome sequence data available, the opportunity to explore predicted effects in specific clinical scenarios is increasing^43^. Unbalanced loss to follow-up by genotype (as a result of death or behaviour) and the complications of employing extensive and complex GRS predictors^15^ (in RbG^mv^) are potentially limiting factors to this approach. Whilst not specific to RbG, these limitations will have an impact on the outcomes of this type of design and will benefit from the study of recruitment dynamics in large-scale prospective studies^45^. As is the case for other forms of MR, these limitations highlight the role of RbG as only part of a required triangulation of evidence when asserting causality or mechanism. As ever, we should not ignore the importance of replication and validation even where studies are targeting the clarification of genetic effects or the impact of suspected causal risk factors.

### Conclusions and recommendations

Considering RbG as a vehicle for undertaking detailed and causal dissection of genetic effects, there are recommendations that come from early experiences with studies of this design:

i. RbG designs are not right for all studies. Dependent on the nature of the genetic variation in question, the sample type or participant recruitment opportunities and the outcomes of interest, there will be optimal conditions for either RbG^sv^ or RbG^mv^ study designs, which should be carefully considered before undertaking such an exercise.
ii. Genetic variant(s) should be independently characterised and, for RbG^mv^, be ideally understood. The genetic variation forming the recall strata are the fundamental building blocks of this study design and the most likely reason that such a study would fail. The integrity of the genetic signal motivating the study should be thoroughly assessed.
iii. The full financial and non-financial costs of undertaking an RbG study should be considered. A deep phenotyping exercise based around an RbG^sv^ design may yield a definitive single hypothesis answer, but the utility of the sampling frame will be limited by that specific study design. This does not render the by-product resources useless by any means (given the randomised nature of their strata), but needs thought.
iv. Transparency, communication (including appropriate disclosure) and thoughtful process in working with participants in RbG studies are paramount. This is a relatively novel approach using genetic data and, whilst the paradigm is simple, the implications are often not.
v. Network RbG studies may be an answer to some of the issues of allele frequency, optimising phenotypic expertise, standardising consent and strategy, and reducing the complexity of original study initiation. Studies do exist capable of RbG and the federated use of these as a network of RbG resources has real potential.

Overall, RbG study designs have the potential to offer independent and informative biological gradients over which specifically designed studies can interrogate the detailed architecture of confirmed associations. In tandem with the driving forces of larger hypothesis-free association studies, the presence of directed follow-up and causal investigation may provide the opportunity to convert some of these outputs into targets for clinical use and future development.

##### BOX 1: Examples of RbG studies

###### Melatonin signalling and Type 2 Diabetes

Several GWASs have identified >100 genetic variants associated with type 2 diabetes (T2D), including a common variant (MAF=0.3) in the melatonin receptor 1b gene (*MTNR1B*). However, the mechanism by which melatonin affects glucose metabolism and development of T2D remains elusive. Tuomi *et al.*^46^ demonstrated that rs10830963, an eQTL for *MTNR1B* in human islets, affects insulin release. To test the hypothesis that activation of *MTNR1B* would result in a reduction of glucose-stimulated insulin secretion, Tuomi *et al.* employed an RbG^sv^ study design. 23 non-diabetic individuals with two copies of the risk allele (GG) and 22 individuals with two copies of the non-risk allele (CC) were recruited for the study where they received 4mg of melatonin for 3 months. The participants underwent an oral glucose tolerance test before and after 3 months of melatonin treatment and levels of plasma glucose, insulin, glucagon and melatonin were measured. The study found that melatonin treatment inhibits insulin secretion, with risk allele carriers exhibiting higher glucose levels. Results from this RbG^sv^ study suggest that melatonin might have a protective role in preventing nocturnal hypoglycemia.

###### *IL2RA* polymorphisms and T cell function

In type 1 diabetes (T1D), the malfunction of CD4^+^ regulatory T cells (Tregs) results in T-cell mediated autoimmune destruction of the pancreatic beta cells. There is a growing body of evidence that interleukin 2 (IL-2) plays a key role in the regulation of Treg function^47-53^. The function of Tregs may be influenced by gene polymorphisms in the IL-2/IL-2 receptor alpha (IL2RA) pathway. Several interleukin-2 (IL-2) receptor alpha-chain (*IL-2RA*) gene haplotypes (rs12722495, rs11594656 and rs2104286) have been shown to be associated with T1D^54^. To investigate if the *IL-2RA* haplotypes are associated with different expression of IL2RA on the surface of peripheral blood T-cells, Dendrou *et al*.^55^ employed an RbG^sv^design, recruiting 50 homozygous or heterozygous individuals for each of the 3 protective haplotypes and 50 homozygous individuals for the susceptible haplotype. Blood samples were collected and the surface expression of IL2RA on peripheral blood T-cells was measured using polychromatic flow cytometry. The study found that individuals with the protective rs12722495 haplotype in *IL-2RA* had increased expression of IL2RA on the surface of memory CD4^+^ T-cells and increased IL-2 secretion compared to individuals with the susceptible haplotypes or those with the protective rs11594656 or rs2104286 haplotype. In addition to this study, Garg *et al*.^56^ employed an RbG^sv^ design recruiting healthy individuals according to their genotype at *IL2RA*-rs12722495 to investigate how polymorphisms in *IL2RA* alter Treg function. Blood samples were taken from 34 healthy individuals and T-cell function was tested. The study found that the T1D-susceptibility *IL2RA* haplotype correlated with diminished Treg function via reduced IL-2 signalling. Findings from the RbG studies by Dendrou *et al.* and Garg *et al.* informed the design of a successful dose-finding, open label, adaptive clinical trial design of Aldesleukin^57^, a recombinant interleukin 2 (IL-2), in participants with T1D to investigate whether Aldesleukin could be potentially used to prevent autoimmune disorders such as T1D by targeting Tregs. The trial found that a single ultra-low dose of Aldeskeukin resulted in early altered trafficking and desensitisation of Tregs, suggesting that Aldeskeukin could be potentially useful to prevent T1D.

###### Body mass index and cardiovascular health in early adulthood

Body mass index (BMI) is a known to influence cardiovascular health in mid-to-late life; however, the nature of this association has not been explored systematically in younger ages. Wade *et al.*^58^ used complementary MR and RbG^mv^ methodologies to estimate the causal effect of BMI on detailed measures of cardiovascular health in a population of young and healthy adults from the Avon Longitudinal Study of Parents and Children (ALSPAC). For the RbG study, 418 individuals were recruited based on a genome-wide GRS predicting variation in BMI (N=191/227 from the lower/upper ∼30% of the continuous genome-wide GRS distribution). The nature of the RbG design allowed more detailed cardiovascular phenotyping, including magnetic resonance imaging (MRI)-derived end-diastolic and systolic volume, ventricular mass, total arterial compliance, systemic vascular resistance, stroke volume and cardiac output, to be undertaken in a targeted sample containing a known biological gradient allowing causal inference. Both MR and RbG analyses indicated a causal role of increased BMI on higher blood pressure and left ventricular mass indexed to height^2^.^7^ (LVMI) in young adults. The RbG results extended this by suggesting a causal role of higher BMI on increasing stroke volume and cardiac output.

## Acknowledgments

This work was supported by the Medical Research Council MC_UU_12013/3.

We also acknowledge Professor John Henderson for providing helpful comments on earlier drafts of the manuscript.

The Cardiovascular Epidemiology Unit at the University of Cambridge is supported by the UK Medical Research Council (G0800270), British Heart Foundation (SP/09/002) and NIHR Cambridge Biomedical Research Centre. DSP is supported by the BHF Cambridge Centre of Excellence (RE/13/6/30180) and the Wellcome Trust (105602/Z/14/Z). C.M.L is supported by the Li Ka Shing Foundation and NIHR Oxford Biomedical Research Centre. The EXCEED study at the University of Leicester has been supported by the Medical Research Council (G0902313) and received partial support from NIHR; the views expressed are those of the authors and not necessarily those of the NHS, the NIHR or the Department of Health. MMcC is a Wellcome Trust Senior Investigator and an NIHR Senior INvestigator. REsearch support relevant to this manuscript comes from Wellcome Trust (090532, 098381, 106130), Medical Research Council (MR/L020149/1) and NIH (R01DK098032; U01DK105535)

*Avon Longitudinal Study of Parents and Children (ALSPAC):* We are extremely grateful to all the families who took part in this study, the midwives for their help in recruiting them, and the whole ALSPAC team, which includes interviewers, computer and laboratory technicians, clerical workers, research scientists, volunteers, managers, receptionists and nurses. ALSPAC mothers were genotyped using the Illumina human660W-quad array at Centre National de Génotypage (CNG) and genotypes were called with Illumina GenomeStudio. The UK Medical Research Council and the Wellcome Trust (Grant ref: 102215/2/13/2) and the University of Bristol provide core support for ALSPAC.

## SUPPLEMENTARY MATERIAL

### Recall-by-Genotype Study Planner: *Methods for power calculation*

#### *RbG*^*sv*^ *– Single variant analysis*

Since a ‘RbG^sv^ study’ design involves the recruitment of specific genotype groups (either major and minor homozygotes or major homozygotes and heterozygotes) resulting in two groups (independent of allele frequency), the power calculation is for a two-sample two-tailed t-test. The equation used was taken from the ‘pwr’ library^S1^ in R^S2^ and is adapted from Cohen^S3^. The non-centrality parameter (NCP, γ) is defined as:

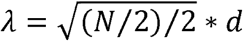

where *N* is the number of individuals in the study (total sample size) and *d* is the standardized effect size. Where the comparison is between minor and major homozygotes, the effect size will be twice the per allele effect at the target locus. Where the comparison is between major homozygotes and heterozygotes, the effect size will be equal to the per allele effect at the target locus.

For a ‘random recall study’ design, where participants are recalled randomly from the population, it is assumed that the sample will contain all three genotypic groups at frequencies determined by the user specified MAF and assuming Hardy-Weinberg equilibrium (HWE). The test of association will therefore manifest as a standard genetic association test. Power is derived from the NCP of a 𝒳^2^-test of association (Sham & Purcell 2014)^S4^ defined as:

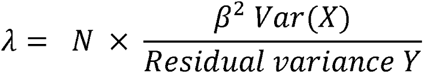

where Y is the trait; X is the allele count at a genetic locus (coded 0, 1 or 2) so that under HWE the variance of *X* (Var(*X*)) is given by 2*p*(1 – *p*), where *p* is the allele frequency at the locus; and β is the regression coefficient of *Y* on *X*. For a minor-effect locus, the residual variance of *Y* is not much smaller than the total variance of 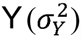, so that the NCP is given by the proportion of trait variance explained by the genetic variant multiplied by the sample size (*N*). Therefore,

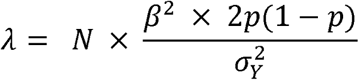

This group represents the comparator, i.e., the power to undertake a nested study of size *N* with no effort to balance genotypic groups through recall.

#### RbG^mv^ – Multiple variant analysis

Power predictions for the ‘RbG^mv^ study’ are determined empirically based on simulated data. Pseudo-individuals from a population cohort (of the size specified by the user) are assigned genotypes at each of the SNPs listed in the GRS variant file according to the effect allele frequency at that SNP. A GRS is generated for each individual either by simply summing the number of risk alleles (unweighted method) or by multiplying the number of risk alleles by their corresponding weight and summing across all SNPs (weighted method).

Exposure phenotypes are simulated by adding a random (normally distributed) error term scaled according to the user-entered r^2^ between the GRS and exposure phenotype. Outcome phenotypes are simulated by adding a random (normally distributed) error term scaled according to the user-entered r^2^ between the exposure and outcome phenotypes to the previously simulated exposure phenotypes. Assuming the random recruitment of individuals from the tails of the GRS distribution (% as specified by user) 1,000 pseudo-datasets are created by selecting N/2 individuals (assuming equal recruitment) from each of the two tails. This procedure is repeated to generate 25 pseudo-populations (each with 1,000 pseudo-datasets).

This simulated data is used firstly to evaluate the power of a ‘RbG^mv^ study’ design to detect a difference in mean exposure phenotype across the strata generated by selecting individuals from the tails of the simulated GRS distribution. A Wilcoxon (Mann-Whitney) test is used to test for a difference in mean exposure phenotype across the two recall groups. The power is estimated as the proportion of test results less than the user specified alpha level across all simulated populations and datasets. This procedure is then repeated to evaluate the power of the same ‘RbG^mv^ study’ design to detect a difference in mean outcome phenotype across the two recall groups.

In addition, we consider the relative power if the same study was performed in either a randomly recruited sample of the same size as the ‘RbG^mv^ study’ (‘random recall study’) or a (genotyped) population cohort of the size specified for recruitment (‘total cohort study’). In both cases, power is derived analytically using the NCP of the 𝒳^2^-test of association between the outcome phenotype and the GRS. The equation used assumes an additive genetic model for a quantitative trait and is presented in Palla and Dudbridge (2015)^S5^ with the NCP defined as:

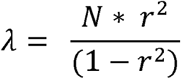

where *N* is the number of individuals (total sample size) and *r*^*^2^*^ is the coefficient of determination between the GRS and either the exposure phenotype or the outcome phenotype; in the case of the latter, the *r*^*^2^*^ between the GRS and the outcome phenotype is calculated as the product of the *r*^*^2^*^ between the GRS and the exposure phenotype and the *r*^*^2^*^ between the exposure phenotype and the outcome phenotype (both provided by the user).

### Data description for Figure 3

The data used to produce Figure 3 in the main text was sourced from mothers recruited by the Avon Longitudinal Study of Parents and Children (ALSPAC). The full details about this cohort can be found below. Physical measures (body mass index (BMI) and systolic blood pressure (SBP)) were recorded at a follow-up clinic carried out approximately 16 years after the mother’s pregnancy. Information about confounding factors was taken from questionnaires completed by the mothers either during pregnancy (education, defined as the mother’s highest qualification) or when their child was aged 18 years (frequency of alcohol consumption and average take-home household income each month). Pre-existing genetic data (for details of the SNP genotyping, imputation, processing and quality control procedures carried out in ALSPAC see below) was used to generate the genetic risk score (GRS) for BMI based on 97 SNPs^S6^. Data analysis was conducted in STATA^S7^ v14.2.

#### ALSPAC: Description of study numbers

ALSPAC recruited 14,541 pregnant women resident in Avon, UK with expected dates of delivery 1st April 1991 to 31st December 1992. 14,541 is the *initial* number of pregnancies for which the mother enrolled in the ALSPAC study and had either returned at least one questionnaire or attended a “Children in Focus” clinic by 19/07/99. Of these *initial* pregnancies, there was a total of 14,676 fetuses, resulting in 14,062 live births and 13,988 children who were alive at 1 year of age.

When the oldest children were approximately 7 years of age, an attempt was made to bolster the initial sample with eligible cases who had failed to join the study originally. As a result, when considering variables collected from the age of seven onwards (and potentially abstracted from obstetric notes) there are data available for more than the 14,541 pregnancies mentioned above.

The number of **new pregnancies** not in the initial sample (known as Phase I enrolment) that are currently represented on the built files and reflecting enrolment status at the age of 18 is 706 (452 and 254 recruited during Phases II and III respectively), resulting in an additional 713 children being enrolled. The phases of enrolment are described in more detail in the cohort profile paper which should be used for referencing purposes: http://ije.oxfordjournals.org/content/early/2012/04/14/ije.dys064.full.pdf. The total sample size for analyses using any data collected after the age of seven is therefore 15,247 pregnancies, resulting in 15,458 fetuses. Of this **total sample** of 15,458 fetuses, 14,775 were **live births** and 14,701 were **alive at 1 year of age**. A 10% sample of the ALSPAC cohort, known as the **Children in Focus (CiF) group**, attended clinics at the University of Bristol at various time intervals between 4 to 61 months of age. The CiF group were chosen at random from the last 6 months of ALSPAC births (1432 families attended at least one clinic). Excluded were those mothers who had moved out of the area or were lost to follow-up and those partaking in another study of infant development in Avon.

#### ALSPAC: Genotyping description

ALSPAC children were genotyped using the Illumina HumanHap550 quad chip genotyping platforms by 23andme subcontracting the Wellcome Trust Sanger Institute, Cambridge, UK and the Laboratory Corporation of America, Burlington, NC, US. The resulting raw genome-wide data were subjected to standard quality control methods. Individuals were excluded on the basis of gender mismatches; minimal or excessive heterozygosity; disproportionate levels of individual missingness (>3%) and insufficient sample replication (IBD < 0.8). Population stratification was assessed by multidimensional scaling analysis and compared with Hapmap II (release 22) European descent (CEU), Han Chinese, Japanese and Yoruba reference populations; all individuals with non-European ancestry were removed. SNPs with a minor allele frequency of < 1%, a call rate of < 95% or evidence for violations of Hardy-Weinberg equilibrium (P < 5E-7) were removed. Cryptic relatedness was measured as proportion of identity by descent (IBD > 0.1). Related subjects that passed all other quality control thresholds were retained during subsequent phasing and imputation. 9,115 subjects and 500,527 SNPs passed these quality control filters.

ALSPAC mothers were genotyped using the Illumina human660W-quad array at Centre National de Génotypage (CNG) and genotypes were called with Illumina GenomeStudio. PLINK^S8^ (v1.07) was used to carry out quality control measures on an initial set of 10,015 subjects and 557,124 directly genotyped SNPs. SNPs were removed if they displayed more than 5% missingness or a Hardy-Weinberg equilibrium P-value of < 1E-6. Additionally SNPs with a minor allele frequency of less than 1% were removed. Samples were excluded if they displayed more than 5% missingness, had indeterminate X chromosome heterozygosity or extreme autosomal heterozygosity. Samples showing evidence of population stratification were identified by multidimensional scaling of genome-wide identity by state pairwise distances using the four HapMap populations as a reference and then excluded. Cryptic relatedness was assessed using a IBD estimate of more than 0.125 which is expected to correspond to roughly 12.5% alleles shared IBD or a relatedness at the first cousin level. Related subjects that passed all other quality control thresholds were retained during subsequent phasing and imputation. 9,048 subjects and 526,688 SNPs passed these quality control filters.

#### ALSPAC: Imputation description

477,482 SNP genotypes in common between the sample of mothers and sample of children were combined. SNPs with genotype missingness above 1% due to poor quality were removed (11,396 SNPs removed). 321 subjects were removed due to potential ID mismatches. This resulted in a dataset of 17,842 subjects containing 6,305 duos and 465,740 SNPs (112 were removed during liftover and 234 were out of HWE after combination). Haplotypes were estimated using ShapeIT (v2.r644) which utilises relatedness during phasing. A phased version of the 1000 genomes reference panel (Phase 1, Version 3) was obtained from the Impute2 reference data repository (phased using ShapeIt v2.r644, haplotype release date Dec 2013). Imputation of the target data was performed using IMPUTE^S9^, ^S10^ V2.2.2 against the reference panel (all polymorphic SNPs excluding singletons), using all 2,186 reference haplotypes (including non-Europeans). This gave 17,842 mothers and children eligible for study with available genotype data. Subsequent consent withdrawals have left 17,825 individuals for study

